# Proteomic profiling of bacterial extracellular vesicles for exploring ovarian cancer biomarkers

**DOI:** 10.1101/2025.04.20.649473

**Authors:** Eri Asano-Inami, Akira Yokoi, Kosuke Yoshida, Kentaro Taki, Masami Kitagawa, Kazuhiro Suzuki, Ryosuke Uekusa, Yukari Nagao, Nobuhisa Yoshikawa, Kaoru Niimi, Yusuke Yamamoto, Hiroaki Kajiyama

## Abstract

Extracellular vesicles (EVs) are present in body fluids and act as disease biomarkers. Emerging evidence has proven that EVs are released not only from mammalian cells but also from bacteria. Ovarian cancer has a dismal prognosis because of difficulties in early detection. This study aimed to identify bacterial EV (BEV) proteins associated with ovarian cancer. Fifteen patients with ovarian cancer or benign tumors were recruited, and EVs were isolated from the ascites. BEVs were recovered from seven in vitro-cultured strains of bacteria present in the vaginal microbiota, and LC-MS/MS analysis was performed. The detected peptide data were annotated to both human and bacterial references, and the human data showed that the profiles of cancer EVs were distinct from those of patients with benign tumors. As analyzed by bacterial proteins, P15636_Protease1 was found as the BEV-associated protein highly expressed in patients with cancer. To distinguish patients with cancer, the area under the curve was 0.88 (95% CI, 0.64–1.00). In addition, homology analysis showed that P15636_Protease1 is a unique protein only detected in bacteria. In this study, ovarian cancer-specific bacterial proteins were identified on EVs, and BEVs in bodily fluids are promising in the discovery of disease biomarkers.

## Introduction

Extracellular vesicles (EVs) are enclosed by lipid bilayers, measure 50–100 nm in size, and are released from all cells. EVs carry various molecules, such as proteins, RNAs, and DNAs, as cargo and can transport them between cells ^1^. EVs are abundant in body fluids, reflect pathological conditions, and have become a focus of attention as cancer-specific diagnostic markers ^2^. In recent years, the association with bacteria has become important in tumor biology. Various tissues and organs possess microbiomes with unique characteristics that contribute to population dynamics and the diversity of microbial species and subspecies ^3 4^. This growing recognition of the importance of the microbiome in health and disease has led to recognition of its influence in the acquisition of cancer malignancy. Pathologists have detected bacteria within solid tumors using advanced profiling techniques. The detection of bacteria within solid tumors has long been recognized by pathologists, and these findings are now being demonstrated using advanced profiling techniques ^4^. Bacteria also release EVs, which were discovered more than 60 years ago in Gram-negative bacteria ^5,6^. Since their discovery in Gram-negative bacteria with outer membranes, they have been recognized as outer membrane vesicles (OMVs) ^7^. However, Gram-positive bacteria also release these vesicles ^8^, which have been called by various names ^6 7^. EVs released by bacteria are referred to as bacterial EVs (BEVs) ^9^. Although pathogens release BEV to deliver toxic determinants to local and distal host cells and shape the immune response ^10,11^, their roles are still vague.

Ovarian cancer is the second most common gynecological malignancy in developed countries and has a poor prognosis and a high mortality rate ^12^. Ovarian cancer has several subtypes, with high-grade serous carcinoma (HGSC) as the most common ^13 14^. However, many uncertainties remain regarding their development, and new molecular targets are required. Vaginal microbiota exists within the vagina ^15^, and the balance in vaginal microbiota can lead to various pathological conditions, such as bacterial vaginosis ^16^, preterm birth ^17^, endometriosis^18^, and ovarian cancer ^19^. A 16s analysis in patients with ovarian cancer showed different ovarian cancer and non-cancer groups have different microbiomes as reflected in ascites ^20^. However, the mechanisms underlying the emergence of these conditions remain unknown, and the platform for clinical applications remains unclear.

In this study, BEVs in patients’ body fluids were analyzed, and they were also recovered from seven strains of bacteria present in the vaginal microbiota. Global proteomic analysis was successfully analyzed. To identify BEV-derived bacterial proteins that are highly expressed in ovarian cancer, ascites samples from 10 patients with ovarian cancer and five without cancer were analyzed. Our findings showed that BEV released by vaginal bacteria can be a potential clinical biomarker for ovarian cancer.

## Methods

### Bacteria culture

*Lactobacillus crispatus* (*L. crispatus*, JCM1185), *Lactobacillus gasseri* (*L. gasseri*, JCM1131), *Lactobacillus jensenii* (*L. jensenii*, JCM15953), *Lismosilactobacillus vaginalis* (*L. vaginalis*, JMC9505) and *Lactobacillus iners* (*L. iners*, JCM12513) were obtained from RIKEN BRC and *Fuso nucleatum* (*F. nucleatum* JNBP_02614) and *Escherichia coli* K-12 (*E. coli*, GTC_2003) were obtained from Gifu University Center for the Conservation of Microbial Genetic Resource, Organization for Research and Community Development, Japan (https://pathogenic-bacteria.nbrp.jp/bacteria/bacteriaAllItemsList.jsp). *L. crispatus, L. gasseri, L. jensenii*, and *L. vaginalis* were first cultured under the aerobic conditions at 37 °C in de Man, Rogosa, and Sharpe (MRS) agar (BD Difco, France). Then, a liquid MRS medium was used for mass culture. *L. iners* was first cultured under anaerobic conditions at 37°C in BL agar (Nissui Pharmaceutical Co, Tokyo, Japan), and a liquid bifidobacterium medium (referring to DSMZ) was then used for mass culture. *F. nucleatum* was first cultured under anaerobic conditions at 37°C in Brucella agar (Kyokutoseiyaku, Tokyo, Japan). We used liquid cultures based on brain heart infusion (Nissui Pharmaceutical Co.) broth medium supplemented with hemin (10 μg/mL, Thermo Fisher Scientific, MA, USA), menadione (5 μg/mL, Nacalai Tesque, Japan), L-cysteine (1 μg/mL, Sigma-Aldrich, MO, USA), and resazurin (2 μg/mL, FUJIFILM Wako Pure Chemical, Japan) as an anaerobic indicator. *E. coli K*-12 was cultured under aerobic conditions at 37°C in L-Broth agar and liquid medium (MP Biomedicals, CA, USA). Bacterial cell concentrations were measured by assessing the optical density at 660 nm (Miniphoto518R, TAITEC), and colony formation units were counted at each OD measurement.

### Isolation of BEVs by differential ultracentrifugation (dUC)

The procedure of separating BEVs used in this study conformed to the standard method of the International Society of Extracellular Vesicles (MISEV2023) ^21^. To obtain an exosome-free medium, the medium for each cell was ultracentrifuged at 32,000 rpm at 4°C overnight. Approximately 400 mL of the medium was cultured with each bacterium for 16–28 h, centrifuged at 2000 × g for 30 min at 4°C to pellet the bacteria, and then centrifuged at 10,000 g 4°C for 40 min (KUBOTA Co., Tokyo, Japan). The pellets were washed with Dulbecco’s phosphate-buffered saline (PBS) and centrifuged again at 10,000 × g for 40 min at 4°C to obtain BEVs. The supernatant was passed through a 0.22-μm filter (Millex-GV 33 mm, Millipore) and ultracentrifuged at 32,000 rpm for 2 h at 4°C. The protein concentrations of EVs and cell lysates were quantified using a Qubit protein assay kit (Thermo Fisher Scientific) with a Qubit 4.0 Fluorometer (Invitrogen Co., MA, USA), according to the manufacturer’s protocol.

### Isolation of BEVs by size exclusion (SEC)

The process was the same as in dUC up to the point where a 0.22-µm filter was used. qEV (iZON Science, Oxford, UK) was used to extract EVs using SEC. The samples were concentrated in an ultracentrifuge to 500 µL. Samples were applied to a qEV column, and a total of 1.6 mL of EVs was eluted according to the protocol. Moreover, 1.6 mL of EV was concentrated to 50 µL using Amicon Ultra 3 K (Merck KGaA, Darmstadt, Germany).

### Nanoparticle tracking analysis (NTA)

The size distribution and particle concentration in EV preparations were analyzed using a NanoSight NS300 nanoparticle tracking analyzer (Malvern Panalytical Ltd., UK). The samples were diluted in PBS and injected at a speed of 100 a. u. into the measuring chamber. The EV flow was recorded in triplicate (30 s each) at room temperature. The equipment settings for data acquisition were kept constant between measurements, with the camera level set at 12.

### Scanning electron microscopy (SEM)

Each bacterial culture was fixed with 2% glutaraldehyde dissolved in PBS at 4°C. The samples were postfixed in 2% osmium tetroxide for 1 h at room temperature and dehydrated using increasing ethanol concentrations (50%–100%). The samples were critical-point dried, stuck on carbon stubs, and coated with osmium (NL-OPC80NS, Japan Laser Corporation, Tokyo, Japan). The samples were imaged under a field-emission electron microscope (JSM-7610F; JEOL Ltd., Tokyo, Japan).

### Transmission electron microscopy (TEM) of BEVs

For TEM, BEVs were prepared by negative staining, EV pellets were resuspended in PBS, and 5 µL of EV samples were loaded on a grid with a carbon support film (Nisshin EM, Tokyo, Japan) and blocked with 1% bovine serum albumin for 1 h at room temperature. After washing with PBS, the EV samples were fixed with 1% glutaraldehyde for 10 min. After washing with distilled water, the EVs were stained with 1% uranyl acetate, and the excess liquid was blotted with filter paper and dried at room temperature. Samples were then examined under a JEM-1400PLUS transmission electron microscope (JEOL Ltd., Tokyo, Japan)

### Patient samples

Patient ascites samples were obtained from female patients admitted at Nagoya University Hospital, Japan. Informed consent was obtained from each patient before surgery. This study was approved by the Ethics Committee of Nagoya University School of Medicine (Approval number: 2017-0053).

Patient details are shown in Table 1.

### Mass Spectrometry

For mass analysis, EV proteins were treated according to Easy Pep™ (Thermo Fisher Scientific) protocols and digested with trypsin for 2 h. The peptides were analyzed by LC-MS using a nanoelectrospray ion source with an Orbitrap Fusion mass spectrometer (Thermo Fisher Scientific) coupled to an UltiMate3000 RSLCnano LC system (Dionex Co., Netherlands) with a nano high-performance LC capillary column (150 mm × 75 μm inside diameter) (Nikkyo Technos Co., Japan). Reversed-phase chromatography was performed with a linear gradient (5%–40% B for 100 min), solvent A (2% acetonitrile and 0.1% formic acid), and solvent B (95% acetonitrile and 0.1% formic acid) at an estimated flow rate of 300 nL/min. Before the MS/MS analysis, a precursor ion scan was performed using a mass-to-charge ratio (*m*/*z*) of 400–1600. MS/MS was performed with an isolation width of 1.6 *m*/*z* with quadrupole, higher-energy collisional dissociation fragmentation with a normalized collision energy of 35%, and rapid scan MS analysis in the ion trap. Only the precursors with charge states 2–6 were sampled for MS2. The dynamic exclusion duration was set to 15 s with 10 parts per million (ppm) tolerance. The instrument was operated in the top-speed mode with 3 s cycles.

### Proteomic data analysis

Raw data were processed for protein identification using Proteome Discoverer (version 1.4, Thermo Fisher Scientific) alone or in conjunction with the MASCOT search engine (version 2.7.0, Matrix Science Inc., Boston, MA, USA). Peptides and proteins were identified with reference to the human protein database or bacterial protein database in UniProt (2021_02), with a precursor mass tolerance of 10 ppm and a fragment ion mass tolerance of 0.8 Da. The fixed modification was set to the carbamidomethylation of cysteine, and the variable modification was set to the oxidation of methionine. Up to two missed tryptic cleavages were permitted

### Homology analysis

Homology analysis was performed using the UniProt basic local alignment search tool (BLAST, https://www.uniprot.org/blast), and UniProtKB Swiss-Prot was selected as the target database to enter the accession number. Homo sapiens and BLAST searches were performed. A sequence with >50% identity was considered homologous. The score is an indicator of sequence similarity that does not depend on the size of the base emitted. E-values <E-4 were considered homologous. The peptide sequences were aligned with the amino acid sequences for comparison with multiple sequence alignment using CLUSTALW (www.genome.jp/tools-bin/clustalw), and the homology was then analyzed using SnapGene (www.snapgene.com).

### Immunoblot analysis

Samples (1 μg) lysed with sEV derived from cancer and non-cancer patients’ ascites were loaded onto polyacrylamide gels and transferred to membranes. After blocking with Blocking One (Nacalai tesque) for 1 h at room temperature, the membranes were incubated overnight at 4 °C with the following primary antibodies: anti-CD9 Antibody, clone MM2/57 CBL162 (Merck, Darmstadt, Germany), CD81 Antibody (B-11) sc-166029 (Santa Cruz Biotechnology, Dallas, TX, USA), and GRP94 Antibody (HL10) sc-393402 (Santa Cruz Biotechnology), which were diluted 1:100 in 10 % Blocking One/Tris-buffered saline with 0.1 % Tween 20 (TBST). The following day, the membranes were washed 3 times for 5 min in TBST, then incubated for 4 h at room temperature with the following secondary antibodies: anti-Mouse IgG, HRP-Linked Whole Ab Sheep NA931 (Cytiva, Tokyo, Japan) were diluted 1:2000 and used for CD9, CD81, and GRP94. Anti-Rabbit IgG. The membranes were imaged using ImageQuant LAS 4010 software (GE Healthcare, Atlanta, GA, USA).

### Statistical analysis

Analyses were conducted using R version 4.0.3 (R Foundation for Statistical Computing, http://R-project.org). The heatmap.2 function of the gplots package (ver. 3.1.0) or Partek Genomics Suite version 7.0 (Partek Inc., MO, USA) was used for the heatmap and hierarchical clustering analysis. To visualize the volcano plots, log2-fold change and adjusted *p*-values for each gene were calculated using the Limma (ver. 1.3.1) and visualized using the EnhancedVolcano (ver1.14.0). The rgl (ver1.3.1) was used for 3D-PCA generation. Triangular correlation plots and receiver operating characteristic (ROC) curves were generated using the corrplot package version 0.92 and pROC package version 1.18.0, respectively. The best combination model for detecting cancer was developed by logistic LASSO regression analysis using the compute.es package version 0.2-5, glmnet package version 4.1-2, hash package version 2.2.6.1, MASS package version 7.3-54, and mutoss package version 0.1-12. The optimal cutoff values for each candidate parameter were set based on the maximum point of the summed sensitivity and specificity (Youden index). The significance for all analyses was defined as *P* < 0.05.

## Results

### Isolation and characterization of EV from human ascites samples

We hypothesized that EVs in ascites from patients with ovarian cancer might contain specific bacterial proteins. EVs were separated from the 15 ascites samples using a qEV column (Figure 1A). NTAs were performed to confirm EV quality. EV particles of approximately 100 nm were observed in both non-cancer and cancer small-EVs (sEVs =30-150 nm in size) (Figure 1B). The presence of sEV was confirmed by TEM, and vesicles approximately 100 nm in size were observed (Figure 1C). The expression of EV-specific proteins was analyzed by immunoblotting. The expression of the EV markers CD81 and CD9 was confirmed, and GRP94, which is not an EV marker, was not observed (Figure 1D).

**Figure 1.**
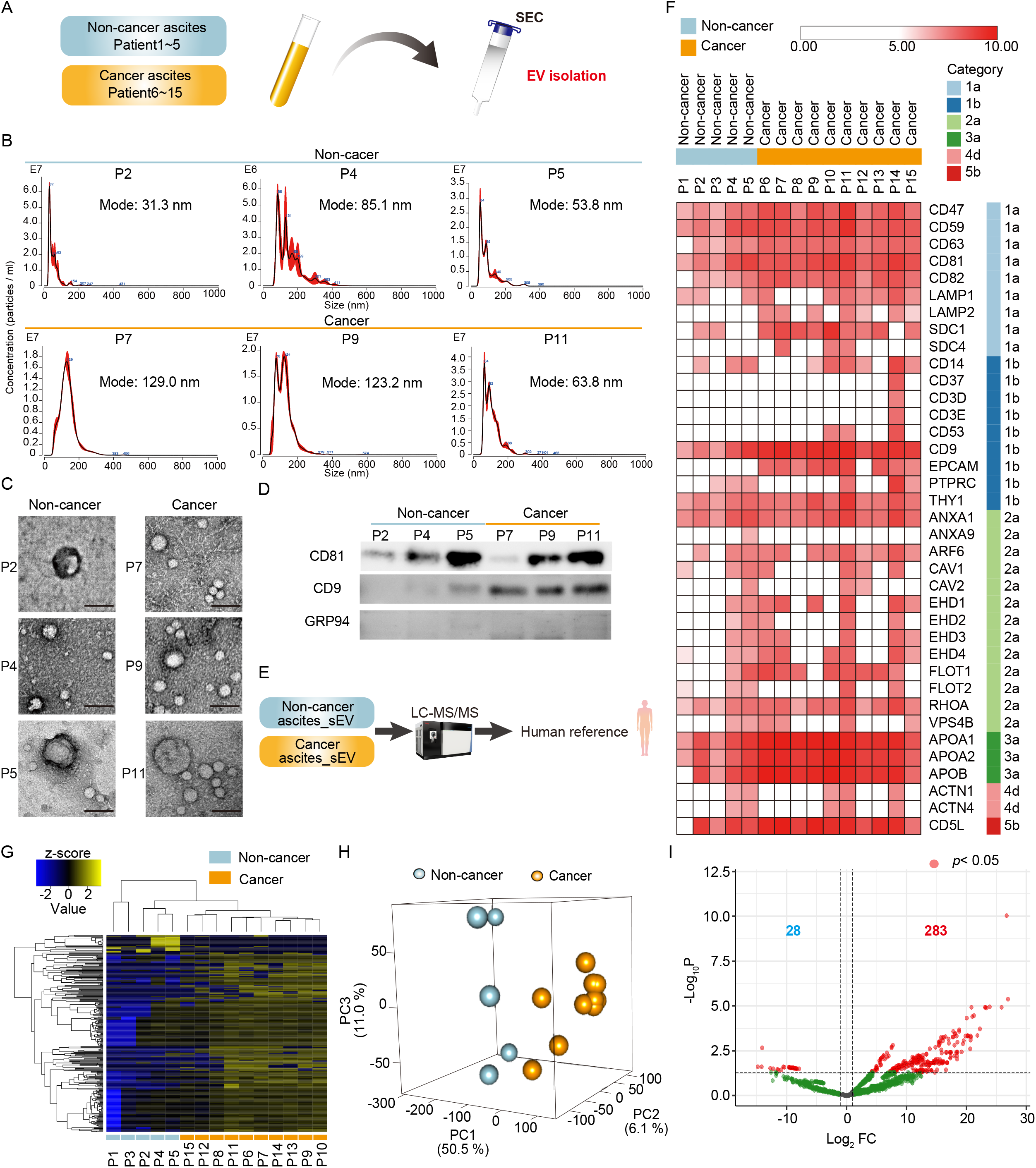
Characterization and mass spectrometry analysis of patient ascites EV. (A) Schematics of EV collection from ascites samples. (B) EVs were recovered from the ascites samples of 10 patients with ovarian cancer and five patients without cancer by size exclusion (SEC). Size distribution obtained by NTAs for isolated EVs derived from representative samples of ascites samples. (C) Transmission electron microscopy analysis of isolated cancer and non-cancer ascites EVs. Scale bar = 100 nm. (D) Immunoblot analyses for CD9, CD81, and GRP94 of EV samples of cancer and non-cancer ascites samples. Uncropped Western blotting data are shown in Figure S1A. (E) Schematics of mass spectrometry analysis of patient ascites EV with human reference. (F) MS data with human reference was obtained for cancer and non-cancer ascites EV. According to protein content–based EV characterization from MISEV2023, EV-associated proteins were categorized into 1a to 5b. Each protein is identified by its gene symbol. The data were converted to a log10 scale. (G) The heat map shows the protein expression obtained from the same MS data as in Figure 1E (total = 2284 proteins). The data were converted to log2 fold change (log2FC). The details of the protein names are given in the Supplementary Information. (H) PCA of the MS data obtained in Figure 1E. (I) Volcano plot analysis between Cancer and non-cancer samples of MS data obtained in Figure 1E. The P-values for each protein were calculated using the Wald test in Limma.

### MS analysis of human ascites EV

Moreover, five non-cancer ascites sEV and 15 cancer ascites sEV were isolated using a qEV column, and MS analysis was performed. The results were analyzed and identified with reference to the human protein database (Figure 1E). Initially, the presence of EV-associated proteins was examined in categories 1–5 of MISEV2023 as a reference. The International Society for Extracellular Vesicles highlighted these categories as indicative of the quality of purified EVs, with the following constituents: transmembrane or glycosylphosphatidylinositol-anchored proteins (category 1), cytosolic proteins (category 2), major constituents of non-EV structures (category 3), larger EV-associated proteins (category 4), and secreted or luminal proteins (category 5). Category 1 proteins were found, and no obvious difference was found between cancer-derived and non-cancer-derived EVs. In other words, sEVs were isolated from samples of patients with cancer to obtain their proteomic profiles (Figure 1F). A heat map drawing of the results of the MS analysis of the human protein reference showed differences between the cancer and non-cancer samples (Figure 1G). The principal component analysis (PCA) also showed differences in profiling between the cancer and non-cancer samples (Figure 1H). Volcano analysis then revealed that in total, 311 of the 2284 proteins were variable in non-cancer and cancer, 283 of them were highly expressed in cancer, and 28 proteins were lowly expressed in cancer (P < 0.05) (Figure 1I).

### Collection of the Vaginal BEV

Moreover, we attempted to recover BEVs from cultured vaginal bacteria and identify the protein. Bacteria that were both endogenous and potential pathogens present in the vagina were selected, referring to the article by Chen et al. ^22^. The optimal culture medium for each bacterium was prepared and cultured. Each bacterium was captured by SEM (Figure 2A). The incubation time to recover EVs from these seven bacteria was then determined. OD values were measured during incubation, and the maximum OD was defined as the point at which the growth of the bacteria reached a plateau. All BEVs were collected approximately 4 h after the bacteria reached their maximum OD. BEVs were collected from seven different bacteria, and the exact BEV recovery was confirmed by TEM and NTA. EVs measuring approximately 100 nm were identified from all bacteria (Figure 2B and 2C).

**Figure 2.**
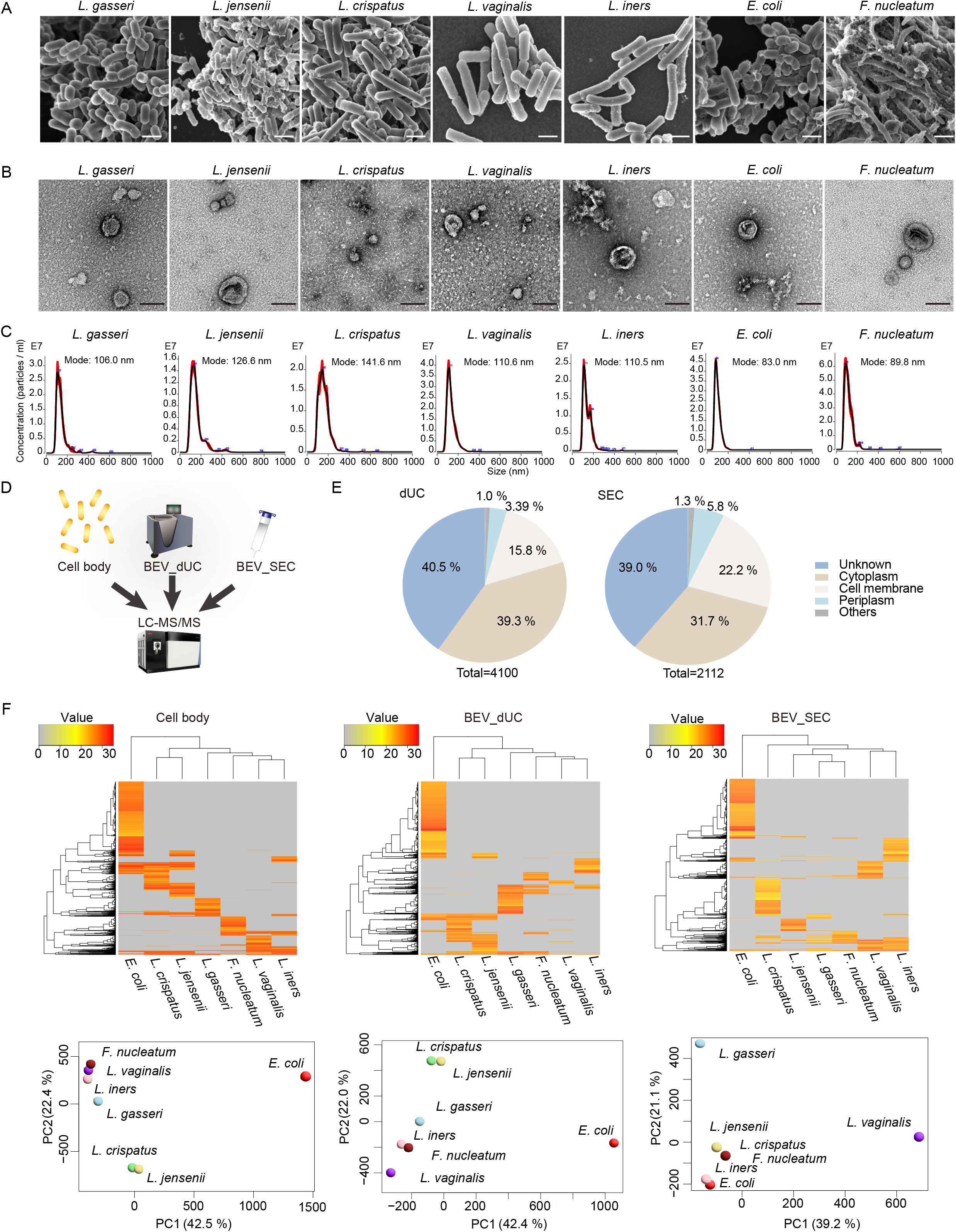
Characterization and mass spectrometry (MS) analysis of the BEVs. (A) Scanning electron microscope images of selected vaginal bacteria. Scale bar = 1 µm (B) Transmission electron microscope images of EVs isolated by size exclusion (SEC) from each bacterium. Scale bar = 100 nm (C) Size distribution obtained by NTAs for isolated EVs derived from representative bacterial samples. (D) Schematics of the mass spectrometry (MS) analysis of BEVs. (E) Circular graph showing the percentage of protein localization obtained by MS of bacterial EVs obtained by dUC and SEC. (F) Heat map and PCA of the expression of MS data obtained from respective samples (cell body = 7659 proteins, BEV_dUC= 4100 proteins, and BEV_SEC = 2112 proteins). The data were converted to log2FC. The details of the protein names are given in the Supplementary file.

### MS analysis of the BEVs

BEVs were then recovered from each of the seven bacteria and each of the fungi by SEC and dUC, respectively, and analyzed by MS (Figure 2D). The proteins identified in each BEV_dUC and BEV_SEC were then classified based on their localization. BEV_dUC identified approximately 40% of the total 4100 proteins, followed by unknown and cytoplasmic proteins (39%) and plasma membrane proteins (16%) (Figure 2E). Conversely, of the 2112 proteins identified by BEV_SEC, approximately 40% were unknown proteins, 31% were cytoplasmic proteins, and 22% were plasma membrane proteins (Figure 2E). As shown in Figure 2F, the protein expression of each bacterium, BEV_dUC, BEV_SEC, or the bacterium itself, was analyzed by MS. The heat map and PCA showed that *E. coli* profiled completely differently in the bacterium; however, some similarities were present in the BEVs, with completely different results for the bacterium itself and EV. *L. crispatus* and *L. jensenii* showed similar profiles for both the bacteria and EVs, whereas the SEC_EVs *L. vaginalis* showed different profiles from the others and changes in the EV recovery methods.

### BEV profiling in ascites samples from patients with ovarian cancer

We attempted to detect EV-derived bacterial proteins that were variably expressed in patients with ovarian cancer. Bacterial proteins were identified from patient-derived EVs by proteomics of EVs recovered from ascites samples of patients with and without cancer (Figure 1E) to a bacterial reference (Figure 3A). As shown in Figure 3B, 212 proteins were successfully detected, and the cancer and non-cancer cell profiles were different in heat map analysis. Cancer and non-cancer profiles were also different in the PCA (Figure 3C). Volcano analysis identified 25 proteins that were significantly expressed in cancers and 10 proteins that were significantly expressed (p < 0.05) (Figure 3D).

**Figure 3.**
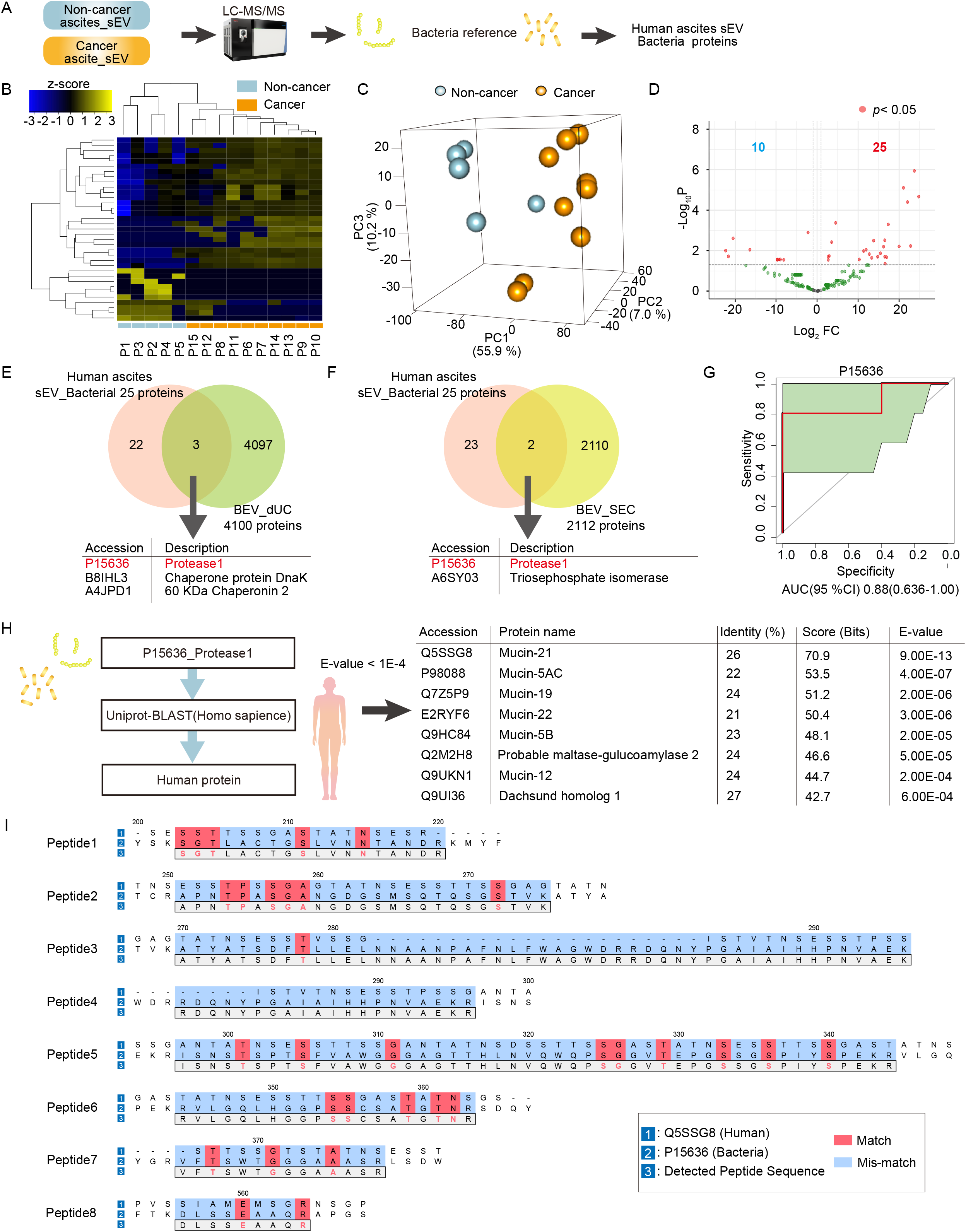
Analysis of bacterial-derived proteins in human ascites EVs. (A) Schematics of the mass spectrometry (MS) analysis of patient ascites EV with bacteria reference. (B) The heat map shows the protein expression obtained from the same MS data as in Figure 3A (total of 35 proteins). The data were converted to log2FC. The details of the protein names are given in the Supplementary Information. (C) PCA of the MS data obtained in Figure 3A. (D) Volcano plot analysis of MS data was obtained in Figure 3A. The P-values for each protein were calculated using the Wald test in Limma. (E) Venn diagram analysis of 25 bacterial proteins highly expressed in cancer obtained from human sample EVs and MS analysis of EVs obtained by dUC from the supernatants of cultured bacteria. (F) The same graph as in Figure 3E for BEVs obtained by SEC (G) ROC-AUC analysis for P15636_Protease1. Values are AUC means (95 % CI). (H) Schematics of the homology analysis of P15636_Protease1. Results of homology analysis of P15636_Protease1. (I) P15636_Protease1 peptide sequences 1-8 were aligned with the amino acid sequence of Q5SSG8_Mucin-21, with matches indicated in red and mismatches in blue. The amino acid number of Q5SSG8 is shown.

Moreover, the Venn diagram was used to search for 25 EV-derived bacterial proteins highly expressed in patients with ovarian cancer and detected in cultured vaginal bacteria-derived EVs. As shown in Figure 3E, 4100 proteins identified in cultured vaginal bacteria-derived bacterial EVs recovered from dUC shared three proteins in the Venn diagram with 25 bacterial proteins from EVs of patients with ovarian cancer. Proteins identified in cultured vaginal bacteria-derived bacterial EVs recovered by dUC and 25 EV bacterial proteins from patients with ovarian cancer (Figure 3E). Venn diagram analysis of the 2112 proteins obtained by BEV_SEC (Figure 2E) revealed two common proteins (Figure 3F). In addition, one protein (Accession no. P15636_Protease1) was shared by both dUC and SEC (Figure 3E and 3F). These proteins were selected as candidate ovarian cancer-associated BEV markers. To evaluate whether P15363_Protease1 is an effective therapeutic target for patients with ovarian cancer, the ROC area under the curve (AUC) analysis was performed. The resulting AUC (95% CI) of P15363_Protease1 was 0.88 (95 % CI: 0.64–1.00), indicating that it is a candidate therapeutic target (Figure 3G). In addition, to P15636_Protease1 (Figure 3G), the B8IHL3_chaperone protein DnaK, A4JPD1_60 kDa chaperonin 2, and A6SY03_ Triosephosphate isomerase (B8IHL3_AUC= 0.96 (95 % CI: 0.87-1.00); A4JPD1_AUC=0.98 (95 % CI: 0.92-1.00); A6SY03_AUC=0.84, (95 % CI: 0.61-1.00) were analyzed (Figure S 1C). LASSO regression analysis revealed improved diagnostic utility (AUC, 95% CI, 0.88–0.96) Model = (−3.118e-11)*P15636 + (5.623e-09)*B8IHL3 + 1.285 (Figure S1D). In summary, our identified BEVs, which are common in vivo and in vitro, are valid therapeutic targets, and the combination of the identified proteins increases their reliability.

### Homology analysis of bacterial proteins

To confirm that P15636_Protase1, which was identified as a potential therapeutic target for ovarian cancer, is a protein of bacterial origin, a homology analysis was performed using the identified peptide sequence. First, UniProt BLAST was run on human CD81 identified from the MS results of human ascites EV, and as expected, human CD81 was 100% homologous (Figure S2A). The four identified CD81 peptides were analyzed using UniProt BLAST and identified as 100% identical to the human protein sequence (Figure S2B). UniProt BLAST analysis of each of the identified CD81 peptides showed 100% homology (Figure S2B and S2C). Homology analysis was performed in the same manner for the bacterial proteins. As shown in Figure 3H, the E-value of P15636_Protease1 was analyzed using UniProt BLAST (*Homo sapiens*) to search for homologous human proteins, and those with E-values less than 1E-4 were considered homologous and listed. All human proteins were <50% homologous, suggesting that P15636_Protease1 was not a human-derived protein but a bacterial protein. Then, the homology of the P15636_Protease1 peptide sequence identified by MS to the Mucin-21 sequence listed at the top of the homology analysis (Figure 3H) was examined. Peptides identified in P15636_Protease1 that had the same sequence (Figure S2D) were excluded. As shown in Figure S2D, the peptide homology was <50%, indicating that the peptide unit was not derived from a human protein but from a bacterial protein. In addition to. P15636_Protease1, homology analysis of the three EV-derived bacterial proteins identified in Figure 3E and 3F showed that some proteins had homology to human proteins, but not the same sequence (= 100% homology), indicating that these proteins are also of bacterial origin. We examined whether the B8IHL3_chaperone protein DnaK, A4JPD1_60 kDa chaperonin 2, and A6SY03_ Triosephosphate isomerase were derived from bacterial proteins in the same way as in Figure 3H. As shown in Figure S2E–2I, some peptides are highly homologous to humans; however, no identical peptide sequences were identified. These proteins were of bacterial origin.

## Discussion

BEVs have been the focus of rapidly expanding attention in recent years, and expectations for new therapeutic markers and other applications are growing ^23^. BEVs secreted by pathogenic bacteria act as an infectious system that delivers toxins to host cells. Although the role and pathogenesis of gut microbiota-derived BEVs have been studied ^24 25^, BEVs released by the vaginal microbiota are less known. Research on BEV is expanding, and databases such as EVpedia (http://evpedia.info) ^26^ compile the proteomics of BEVs. Lipopolysaccharide in Gram-negative bacteria and lipoteichoic acid in Gram-positive bacteria are BEV markers ^27 28^. However, common BEV markers were fewer than in mammalian cells, and BEV research is currently limited. In the present study, BEVs were recovered, and proteomics was performed from seven fungi by dUC and SEC. A total of 4100 proteins were identified by dUC and 2112 by SEC. Figure S1B shows that dUC and SEC contained 1491 proteins. This result indicates that dUC may be more efficient in EV recovery or the EVs obtained by dUC may contain many proteins not derived from EVs. Even with EV recovery from mammalian cells, which has been extensively studied than from bacteria, which method is better, such as dUC or SEC, is still being debated ^29 30 31^. As shown in Figure 2F, many bacterial proteins have unknown localization; thus, further analysis is needed to confirm which recovery method is better.

Ovarian cancer is a gynecological cancer with a high fatality rate when metastasis is detected ^32 12 33 34^. Ovarian cancer has various subtypes, most notably HGSC, which accounts for approximately 75% of cases and has a lethality rate of nearly 90% ^35^. However, no specific and sensitive biomarker has been established for HGSC, and CA125, the most commonly used biomarker in clinical diagnosis, has proven ineffective in the early detection of ovarian cancer ^36^. Therefore, ovarian cancer requires a new marker for its early detection and diagnosis. The vaginal microbiota is commonly associated with alterations in immune and metabolic signaling and the etiology of many gynecologic events such as vaginitis, pelvic inflammatory disease, endometritis, and gynecologic cancers ^37^. Several studies have reported alterations in the vaginal microbiota of patients with ovarian cancer ^38 39^. The ovaries are located in the abdominal cavity, and ascites reflect the pathogenesis of ovarian cancer ^40^. Despite reports of bacterial changes in ascites fluid 16S-seq from patients with ovarian cancer and without cancer ^20^, no studies have reported on bacteria-derived proteins in EVs.

In this study, a bacterial protein, P15636_Protease1, differentially expressed in patients with and without cancer was identified as a potential candidate for therapeutic prediction (Figure 3G). Homology analysis was performed by BLAST to confirm the bacterial proteins. No homologous human proteins were found, indicating that the peptides identified were of bacterial origin (Figure 3I). Homology analysis using BLAST ^41 42^ is generally popular; however, various other homology analysis tools, such as FASTA ^43^, are available, and which method is better is still being debated ^44^. Therefore, future studies are needed to examine whether proteins such as P15636_Protease1 are expressed in EVs derived from the vaginal microbiota, and more comprehensive patient samples are necessary to generate specific antibodies and further investigate the expression of P15636_Protease1. There have been no cases in which bacterial proteins were used as markers or predictors of treatment in ovarian cancer; thus, functional analysis of this protein may lead to new approaches to ovarian cancer diagnosis.

In summary, this study identified a bacterial protein that is highly expressed in the ascites EVs of patients with ovarian cancer and showed that the protein is a candidate marker for ovarian cancer treatment. Currently, no bacterial protein markers for ovarian cancer have been established. This study provides a new potential marker and therapeutic target for ovarian cancer treatment, which may lead to the early detection of ovarian cancer and improvement of prognosis.

## Abbreviations

*EV*: Extracellular vesicle
*BEV*: Bacterial Extracellular vesicle
*OMV*: Outer membrane vesicle
*HGSC*: High-grade serous carcinoma
*MISEV*: The standard method of the International Society of Extracellular Vesicles
*dUC*: Differential ultracentrifugation
*SEC*: Size exclusion
*SEM*: Scanning electron microscopy
*TEM*: Transmission electron microscopy
*NTA*: Nanoparticle tracking analysis
*AUC*: Area under the curve
*ROC*: Receiver operating characteristic
*PCA*: Principal component analysis
*LC-MS*: Liquid chromatograph-mass spectrometer
*CA125*: Cancer antigen 125
*BLAST*: Basic Local Alignment Search Tool
*UniProt*: The Universal Protein Resource
*FASTA*: Fast-all
*L. crispatus*: *Lactobacillus crispatus*
*L. gasseri*: *Lactobacillus gasseri*
*L. jensenii*: *Lactobacillus jensenii*
*L. vaginalis*: *Lismosilactobacillus vaginalis*
*L. iners*: *Lactobacillus iners*
*F. nucleatum*: *Fuso nucleatum*
*E. coli*: *Escherichia coli*

## Acknowledgments

We thank the Division for Medical Research Engineering, Nagoya University Graduate School of Medicine, for equipment usage, technical support, and so on.

This work was financially supported by the following grants: the Practical Research for Innovative Cancer Control, Japan Agency for Medical Research and Development (AMED) grant numbers 24ck0106951h0001; the Fusion Oriented Research for disruptive Science and Technology (FOREST; JPMJFR204J) from Japan Science and Technology Agency (JST); JSPS KAKENHI grant numbers 23KJ1136, 24K12528 and 24K02586. Moreover, The Naito Foundation, and Tokai Pathways to Global Excellence (T-GEx) as part of NEXT Strategic Professional Development Program for Young Researchers, also supported this work.

## Author contributions

E.A-I. and A.Y., conceived the idea, designed the experiments, and interpreted the results. Each author contributed to this work as follows: E.A.-I., isolated EVs and performed NTA and WB; E.A.-I. cultured bacteria; E.A.-I. and M.K performed TEM; E.A.-I. and K.T., performed LC-MS/MS; K.Y and Y.Y., analyzed proteomics results; R.U, N.Y, M.Y, S.T, N.Y, and K.N., clinical sample collection; K.S analyze ROC curve; H.K., supervising clinical concepts; E.A.-I. performed homology analysis; E.A-I. and A.Y., wrote the manuscript, and all authors critically reviewed the paper.

## Ethics approval statement

The study was approved by the Institutional Ethics Board of Nagoya University (approval number: 2017□0497). Pre□existing samples and medical records were used. Thus, we provided disclosure information on the methods of this study and gave the subjects opportunities to reject enroll in this study.

## Patient consent statement

Written informed consent was obtained from all patients, and all participants agreed to publication.

## Data and materials availability

All data needed to evaluate the conclusions in the paper are present in the paper and/or the Supplementary Materials. Additional data related to this paper is available from the authors on reasonable request.

The proteomics LC-MS/MS raw data results have been deposited to the MassIVE ^45^ partner repository with the dataset identifier: MSV00009719

## Reference

1. Yokoi A., Ochiya T. Exosomes and extracellular vesicles: Rethinking the essential values in cancer biology. Semin Cancer Biol 74, 79–91 (2021).

2. Xu R., Rai A., Chen M., Suwakulsiri W., Greening D. W., Simpson R. J. Extracellular vesicles in cancer - implications for future improvements in cancer care. Nat Rev Clin Oncol 15, 617–638 (2018).

3. Hanahan D. Hallmarks of Cancer: New Dimensions. Cancer Discov 12, 31–46 (2022).

4. Nejman D., et al. The human tumor microbiome is composed of tumor type-specific intracellular bacteria. Science 368, 973–980 (2020).

5. Chatterjee S. N., Das J. Electron microscopic observations on the excretion of cell-wall material by Vibrio cholerae. J Gen Microbiol 49, 1–11 (1967).

6. Devoe I. W., Gilchrist J. E. Release of endotoxin in the form of cell wall blebs during in vitro growth of Neisseria meningitidis. J Exp Med 138, 1156–1167 (1973).

7. Jan A. T. Outer Membrane Vesicles (OMVs) of Gram-negative Bacteria: A Perspective Update. Front Microbiol 8, 1053 (2017).

8. Toyofuku M., Nomura N., Eberl L. Types and origins of bacterial membrane vesicles. Nat Rev Microbiol 17, 13–24 (2019).

9. Tulkens J., De Wever O., Hendrix A. Analyzing bacterial extracellular vesicles in human body fluids by orthogonal biophysical separation and biochemical characterization. Nat Protoc 15, 40–67 (2020).

10. Ellis T. N., Kuehn M. J. Virulence and immunomodulatory roles of bacterial outer membrane vesicles. Microbiol Mol Biol Rev 74, 81–94 (2010).

11. Kaparakis-Liaskos M., Ferrero R. L. Immune modulation by bacterial outer membrane vesicles. Nat Rev Immunol 15, 375–387 (2015).

12. Torre L. A., Bray F., Siegel R. L., Ferlay J., Lortet-Tieulent J., Jemal A. Global cancer statistics, 2012. CA Cancer J Clin 65, 87–108 (2015).

13. Kossaï M., Leary A., Scoazec J. Y., Genestie C. Ovarian Cancer: A Heterogeneous Disease. Pathobiology 85, 41–49 (2018).

14. Shih I. M., Wang Y., Wang T. L. The Origin of Ovarian Cancer Species and Precancerous Landscape. Am J Pathol 191, 26–39 (2021).

15. Moreno I., et al. Evidence that the endometrial microbiota has an effect on implantation success or failure. Am J Obstet Gynecol 215, 684–703 (2016).

16. Onderdonk A. B., Delaney M. L., Fichorova R. N. The Human Microbiome during Bacterial Vaginosis. Clin Microbiol Rev 29, 223–238 (2016).

17. Peelen M. J., et al. The influence of the vaginal microbiota on preterm birth: A systematic review and recommendations for a minimum dataset for future research. Placenta 79, 30–39 (2019).

18. Muraoka A., et al. Fusobacterium infection facilitates the development of endometriosis through the phenotypic transition of endometrial fibroblasts. Sci Transl Med 15, eadd1531 (2023).

19. Zhao X., Liu Z., Chen T. Potential Role of Vaginal Microbiota in Ovarian Cancer Carcinogenesis, Progression and Treatment. Pharmaceutics 15, (2023).

20. Yu B., et al. Identification of fallopian tube microbiota and its association with ovarian cancer. Elife 12, (2024).

21. Welsh J. A., et al. Minimal information for studies of extracellular vesicles (MISEV2023): From basic to advanced approaches. J Extracell Vesicles 13, e12404 (2024).

22. Chen C., et al. The microbiota continuum along the female reproductive tract and its relation to uterine-related diseases. Nat Commun 8, 875 (2017).

23. Gurunathan S., Kim J. H. Bacterial extracellular vesicles: Emerging nanoplatforms for biomedical applications. Microb Pathog 183, 106308 (2023).

24. Peregrino E. S., et al. The Role of Bacterial Extracellular Vesicles in the Immune Response to Pathogens, and Therapeutic Opportunities. Int J Mol Sci 25, (2024).

25. Liang A., Korani L., Yeung C. L. S., Tey S. K., Yam J. W. P. The emerging role of bacterial extracellular vesicles in human cancers. J Extracell Vesicles 13, e12521 (2024).

26. Kim D. K., et al. EVpedia: an integrated database of high-throughput data for systemic analyses of extracellular vesicles. J Extracell Vesicles 2, (2013).

27. Tulkens J., et al. Increased levels of systemic LPS-positive bacterial extracellular vesicles in patients with intestinal barrier dysfunction. Gut 69, 191–193 (2020).

28. Champagne-Jorgensen K., Mian M. F., McVey Neufeld K.A., Stanisz A. M., Bienenstock J. Membrane vesicles of Lacticaseibacillus rhamnosus JB-1 contain immunomodulatory lipoteichoic acid and are endocytosed by intestinal epithelial cells. Sci Rep 11, 13756 (2021).

29. Brennan K., et al. A comparison of methods for the isolation and separation of extracellular vesicles from protein and lipid particles in human serum. Sci Rep 10, 1039 (2020).

30. Takov K., Yellon D. M., Davidson S. M. Comparison of small extracellular vesicles isolated from plasma by ultracentrifugation or size-exclusion chromatography: yield, purity and functional potential. J Extracell Vesicles 8, 1560809 (2019).

31. Gámez-Valero A., Monguió-Tortajada M., Carreras-Planella L., Franquesa M., Beyer K., Borràs F.E. Size-Exclusion Chromatography-based isolation minimally alters Extracellular Vesicles’ characteristics compared to precipitating agents. Sci Rep 6, 33641 (2016).

32. Coleman R. L., Monk B. J., Sood A. K., Herzog T. J. Latest research and treatment of advanced-stage epithelial ovarian cancer. Nat Rev Clin Oncol 10, 211–224 (2013).

33. Webb P. M., Jordan S. J. Epidemiology of epithelial ovarian cancer. Best Pract Res Clin Obstet Gynaecol 41, 3–14 (2017).

34. Goff B. A., Mandel L., Muntz H. G., Melancon C. H. Ovarian carcinoma diagnosis. Cancer 89, 2068–2075 (2000).

35. Integrated genomic analyses of ovarian carcinoma. Nature 474, 609–615 (2011).

36. Menon U., et al. Ovarian cancer population screening and mortality after long-term follow-up in the UK Collaborative Trial of Ovarian Cancer Screening (UKCTOCS): a randomised controlled trial. Lancet 397, 2182–2193 (2021).

37. Łaniewski P., Ilhan Z. E., Herbst-Kralovetz M. M. The microbiome and gynaecological cancer development, prevention and therapy. Nat Rev Urol 17, 232–250 (2020).

38. Nené N.R., et al. Association between the cervicovaginal microbiome, BRCA1 mutation status, and risk of ovarian cancer: a case-control study. Lancet Oncol 20, 1171–1182 (2019).

39. Jacobson D., et al. Shifts in gut and vaginal microbiomes are associated with cancer recurrence time in women with ovarian cancer. PeerJ 9, e11574 (2021).

40. Ford C. E., Werner B., Hacker N. F., Warton K. The untapped potential of ascites in ovarian cancer research and treatment. Br J Cancer 123, 9–16 (2020).

41. Altschul S. F., et al. Gapped BLAST and PSI-BLAST: a new generation of protein database search programs. Nucleic Acids Res 25, 3389–3402 (1997).

42. Zaru R., Orchard S. UniProt Tools: BLAST, Align, Peptide Search, and ID Mapping. Curr Protoc 3, e697 (2023).

43. Altschul S. F., Gish W., Miller W., Myers E. W., Lipman D. J. Basic local alignment search tool. J Mol Biol 215, 403–410 (1990).

44. Webber C., Barton G. J. Increased coverage obtained by combination of methods for protein sequence database searching. Bioinformatics 19, 1397–1403 (2003).

45. Choi M., et al. MassIVE.quant: a community resource of quantitative mass spectrometry-based proteomics datasets. Nat Methods 17, 981–984 (2020).

