## Supplementary material for "Proteomic profiling of bacterial extracellular vesicles for exploring ovarian cancer biomarkers": Supplemenatal Figure

**Supplementary Figure 1**

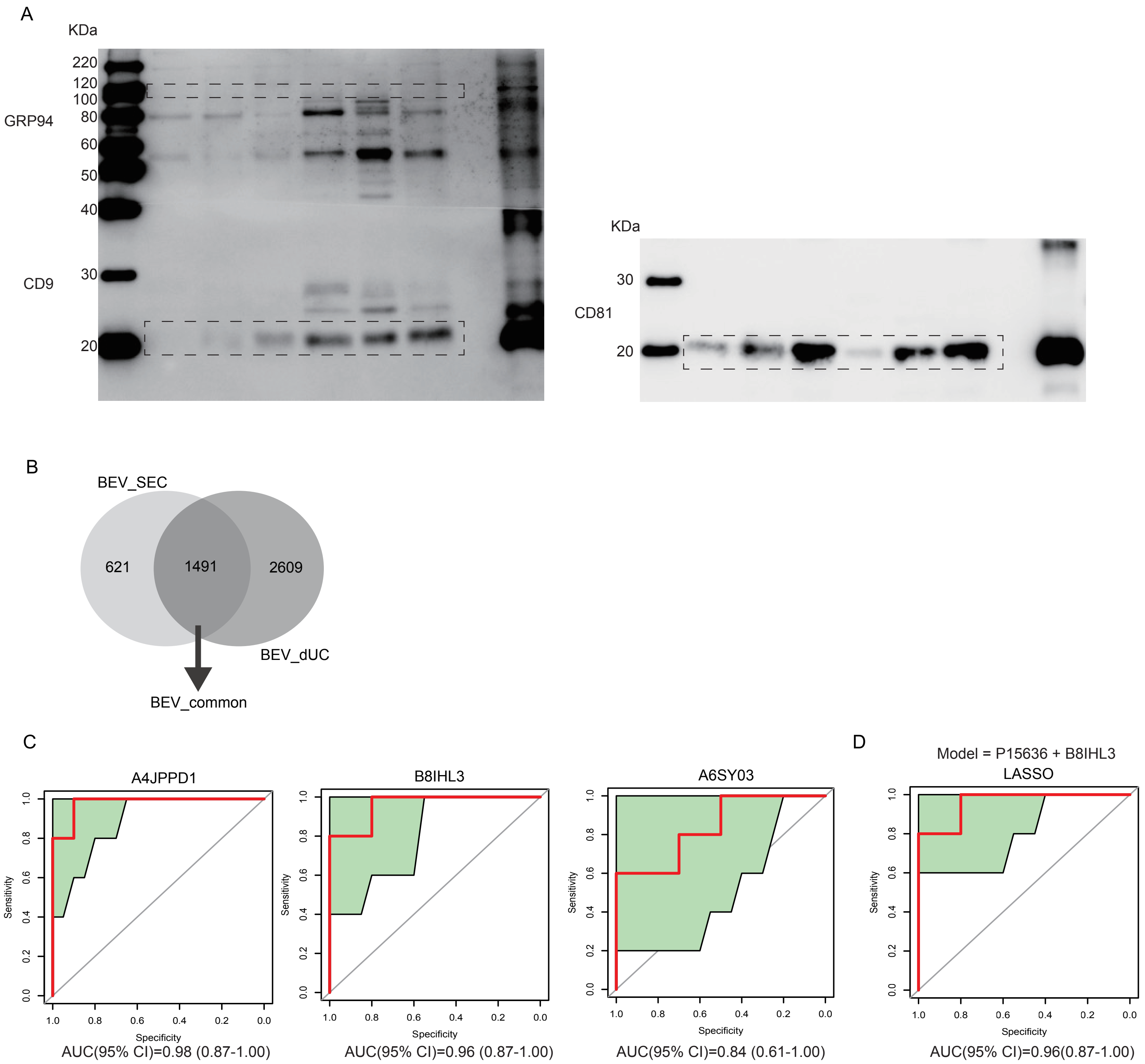

### Supplementary Figure 2

A

| Accession | Protein name | Identity (%) | Score (Bits) | E-value |
|---|---|---|---|---|
| P60033 | CD81 antigen | 100 | 1237 | 7.90E-175 |

B

| CD81_Peptide | Sequence | Identity (%) |
|---|---|---|
| Peptide1 | QFYDQALQQAVVDDDANNAK | 100 |
| Peptide2 | TFHETLDCCGSSTLTALTTSVLK | 100 |
| Peptide3 | NNLCPSGSNIISNLFKEDCHQK | 100 |
| Peptide4 | IDDLFSGK | 100 |

C

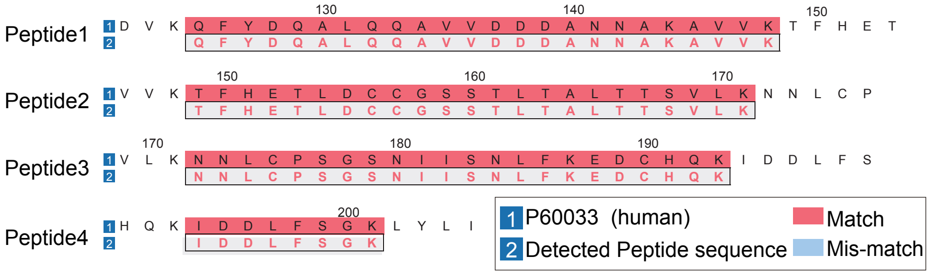


D

| P15636_Peptide | Sequence | Identity (%) |
|---|---|---|
| Peptide1 | SGTLACTGSLVNNTANDRK | 22.2 |
| Peptide2 | APNTPASGANGDGSMSQTWSGSTVK | 24.0 |
| Peptide3 | ATYATSDFTLLELNNAANPAFNLFWAGQDRRDQNYPGAIAIHHPNVAFK | 4.8 |
| Peptide4 | RDQNYPGAIAIHHPNVAEKR | 0 |
| Peptide5 | RISNSTSPTSFVAWGGGAGTTHLNCQWQPSGGVTEPGSSGSPIYSPEKR | 20.8 |
| Peptide6 | SDQYGRVFTSWTGGGAAASR | 25.0 |
| Peptide7 | VFTSWTGGGAAASR | 21.4 |
| Peptide8 | DLSSEAAQR | 22.2 |

E

| Bacteria | Human | Identity (%) |
|---|---|---|
| B8IHL3_DNAK | P38646_GRP75 | 67.2 |
| A4JPD1_CH602 | P10809_CH60 | 53.8 |
| A6SY03_TPIS | P60174_TPIS | 44.5 |

F

| B8IHL3 Peptide | Sequence | Identity (%) |
|---|---|---|
| Peptide1 | QAVTNPSR | 14.3 |
| Peptide2 | IINEPTAAALAYGLDKR | 58.8 |
| Peptide3 | EQGIDLTK | 12.5 |
| Peptide4 | ARTTDLMQASMK | 0 |

G

| A6SY03 Peptide | Sequence | Identity (%) |
|---|---|---|
| Peptide1 | IVVAYEPVWAIGTGK | 86.7 |
| Peptide2 | TATPAMAQEVHAMLR | 73.3 |
| Peptide3 | NAEAAMQVR | 30 |

H

| A4JPD1 Peptide | Sequence | Identity (%) |
|---|---|---|
| Peptide1 | AVGVVLEELR | 50 |
| Peptide2 | AALTDLKGANGDQDAGIR | 50 |

I

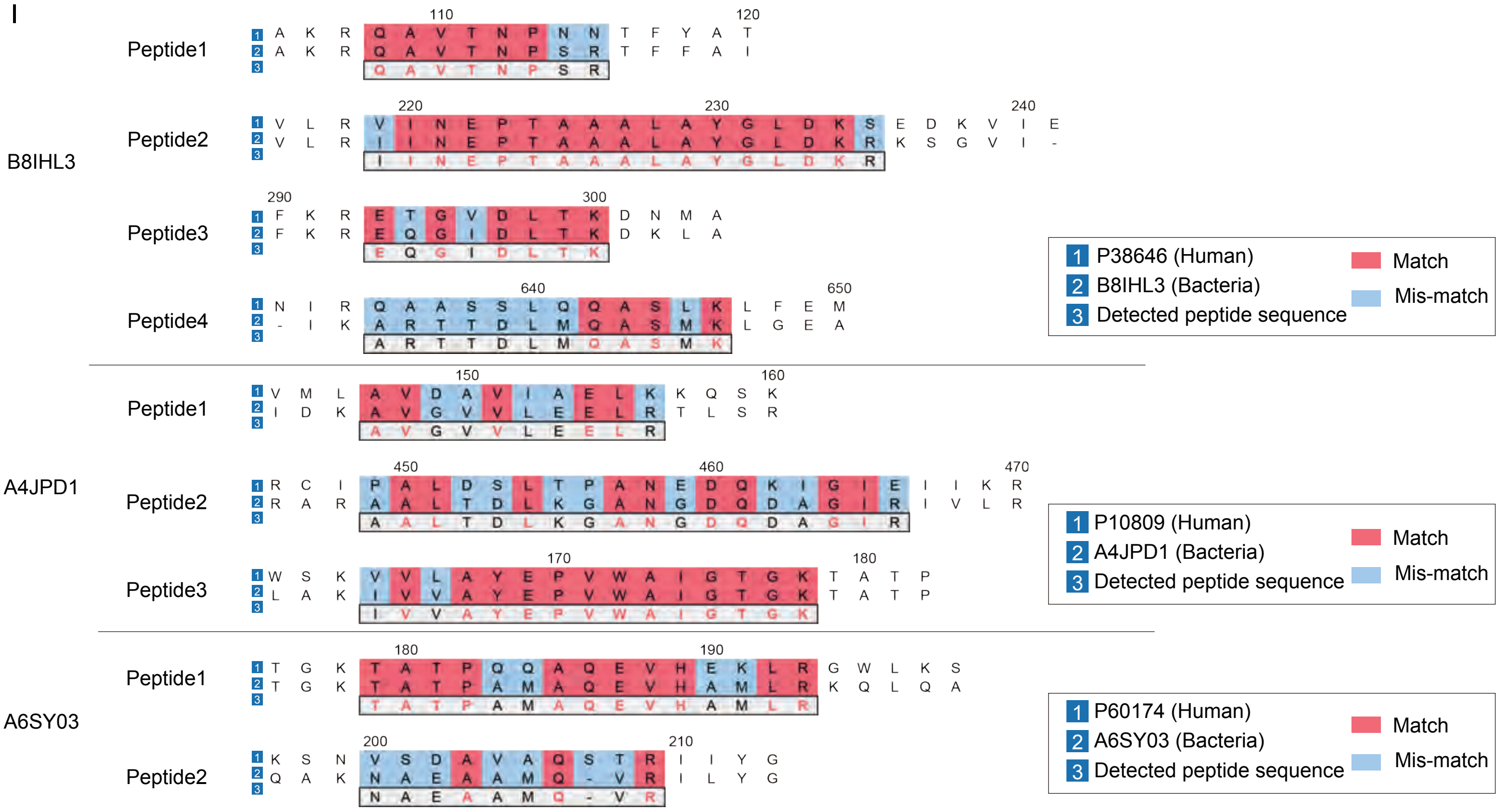
