## Supplemental Figure legends for "Proteomic profiling of bacterial extracellular vesicles for exploring ovarian cancer biomarkers"

**Figure S1**

(A) Uncropped Western blotting images. The membrane was cut between 40 and 50 kDa and detected with GRP antibody on the upper side and CD9 or CD81 antibody on the lower side. (B) Venn diagram analysis of the mass spectrometry analysis of BEVs obtained by dUC or SEC from the supernatants of cultured bacteria. (C) ROC analysis for B8IHL3_chaperone protein DnaK, A4JPD1_60 kDa chaperonin 2, and A6SY03_ Triosephosphate isomerase. Values are AUC means (95% CI). (D) LASSO analysis of P15363 and B8IHL3. Values are AUC means (95% CI).

**Figure S2**

(A) Human CD81 sequence analyzed by UniProt BLAST. (B) The peptide sequences 1–4 of CD81 identified by MS in Figure 1E were aligned to the protein sequence of CD81 human for homology analysis. (C) Aligning peptide sequences of CD81 with the amino acid sequence of P60033_CD81, with matches in red and mismatches in blue. The amino acid number of P60033 is shown. (D) Homology analysis of peptide sequences 1–8 of P15636_Protease1 to Q5SSG8_Mucin-21. (E) Results of the homology analysis of B8IHL3_chaperone protein DnaK, A4JPD1_60 kDa chaperonin 2, and A6SY03_ Triosephosphate isomerase. (F) to (H) Homology analysis of the peptide sequence of the B8IHL3_chaperone protein DnaK to P38646_GRP75 (F), A4JPD1_60 kDa chaperonin 2 to P10809_CH60 (G), or A6SY03_ Triosephosphate isomerase to P60174_TPIS (H). (I) Aligning the peptide sequences of the B8IHL3_chaperone protein DnaK, A4JPD1_60 kDa chaperonin 2, or A6SY03_ Triosephosphate isomerase with the amino acid sequence of each indicated human protein, with matches in red and mismatches in blue. The amino acid number of P38646, P10809, or P60174 is shown.
