## Supplementary material for "Proteomic profiling of bacterial extracellular vesicles for exploring ovarian cancer biomarkers": Table1

| No. | Sample | Pathology |
| --- | --- | --- |
| 1 | Ascites | seous cystadenoma |
| 2 | Ascites | fibrothecoma |
| 3 | Ascites | fibroma |
| 4 | Ascites | mature teratoma |
| 5 | Ascites | fibroma |
| 6 | Ascites | HGSC |
| 7 | Ascites | HGSC |
| 8 | Ascites | HGSC |
| 9 | Ascites | HGSC |
| 10 | Ascites | HGSC |
| 11 | Ascites | HGSC |
| 12 | Ascites | HGSC |
| 13 | Ascites | HGSC |
| 14 | Ascites | HGSC |
| 15 | Ascites | HGSC |

Table S1
